# Use of untargeted magnetic beads to capture *Mycobacterium smegmatis* and *Mycobacterium avium paratuberculosis* prior detection by mycobacteriophage D29 and Real-Time-PCR

**DOI:** 10.1101/2021.06.25.449992

**Authors:** Gabriel Rojas-Ponce, Dominic Sauvageau, Roger Zemp, Herman W. Barkema, Stephane Evoy

## Abstract

Untargeted magnetic beads were evaluated to capture *Mycobacterium smegmatis* and *Mycobacterium avium* subspecies *paratuberculosis* (MAP) from spiked feces, milk, and urine. Untargeted magnetic beads recovered more *M. smegmatis* cells from PBS suspension than the typical centrifugation method; 96.31% of 1.68 × 10^4^ CFU/mL viable *M. smegmatis* were recovered by beads and 0% by centrifugation. Likewise, the *F57*-qPCR detection of MAP cells was different whether they were recovered by beads or centrifugation; cycle threshold (Ct) was lower (p<0.05) for the detection of MAP cells recovered by beads than centrifugation, indicative of higher sensitivity. Magnetic separation of MAP cells from milk, urine, and feces specimens were detected by *F57* and *IS900* sequences. Ct values demonstrated that beads captured no less than 10^9^ CFU/mL from feces and no less than 10^4^ CFU/mL from milk and urine suspensions. In another detection strategy, *M. smegmatis* coupled to magnetic beads were infected by mycobacteriophage D29. Plaque forming units were observed after 24 h of incubation from urine samples containing 2 × 10^5^ and 2 × 10^3^ CFU/mL *M. smegmatis*. The results of this study provide a promising tool for diagnosis of Johne’s disease and other mycobacterial diseases such as tuberculosis.

## 1. Introduction

The genus *Mycobacterium* is grouped in *Mycobacterium tuberculosis* complex (MTBC) and non-tuberculosis mycobacteria (NTM) (1) and can cause multiple diseases in humans and animals (2,3). *M. tuberculosis* (MTB), the etiological agent of tuberculosis (TB) in humans causes millions of deaths every year, mainly in developing countries (4), can be isolated from sputum (5), lavage fluid (6,7), gastric lavage (8,9), urine (10–12), and stools (13,14). *M. avium* subspecies *paratuberculosis* (MAP), a NTM that cause Johne’s disease (JD) or paratuberculosis, has been isolated from intestinal samples of domestic animals (e.g., cattle, sheep, goats, and farmed deer), wildlife (e.g., rabbit, hare, fox, stoat, weasel, crow, rook); and human beings (15)(16)(17).

Definitive diagnostic of TB and JD is based on the isolation of MTB and MAP from human and animal’s specimens on culture media. For that, the specimen, which can be contaminated with normal flora organisms, should be decontaminated and centrifuged (18). Classical decontamination methods to isolate MTB and MAP using the culture method are based on the use of *N*-acetyl-L-cysteine–sodium hydroxide (NALC-NaOH) (18) and hexadecylpyridinium chloride (HPC) (19), respectively for MTB and MAP, and incorporate the concentration of the specimen by centrifugation at 3,000 *x g* at 4°C for 15-20 min (18–20). Given that these chemical decontaminants can kill a portion of mycobacteria present in the specimen (19–22), and that the availability of refrigerated centrifuges with sealed safety cups and aerosol-tight rotors may be challenging in laboratories from low- and middle-income countries (23), additional efforts should be focused on the development of new and simple methods that enable the separation of MTB and MAP from specimens without affecting their viability to grow on culture media or be detected.

Magnetic-separation methods, already established as routine methods to detect *Listeria monocytogenes* (24), *Salmonella* spp. (25), and *Escherichia coli* O157:H7 (26) (27) in food and veterinary clinical samples, are promising alternatives to selectively separate and concentrate MTB (28–32) and MAP (33,34). Magnetic bead characteristics (composition, size, concentration, and surface modification) and the nature of coating ligands (antibodies, peptides, chemicals) are important factors enabling the concentration of the target bacteria and increasing the performance of further complementary detection methods such as culture, PCR, microscopy, antigen detection immunoassay, or phage assay (33,35,36).

Tosyl-activated magnetic beads (e.g., Dynabeads M-280 Tosylactivated) can bind covalently to primary amine and sulfhydryl groups in proteins and peptides (37). Importantly, some surface proteins have been identified on mycobacteria (38–40). As per their characteristics, we postulate that Tosyl-activated magnetic beads may interact with these surface proteins enabling the capture of *M. smegmatis* and MAP cells in suspension. Because the use of untargeted beads to capture MAP cells would diminish laboratory work and costs, and would increase affordability in countries with a high burden of TB and JD, this study determined the potential of Tosyl-activated Dynabeads M-280 (untargeted magnetic beads) to capture *M. smegmatis* and MAP from spiked urine, feces, and milk samples prior to detection by real-time PCR (qPCR) and phage D29.

## 2. Material and Methods

### 2.1. Bacterial cultures

MAP ATCC 19698 was cultured in Middlebrook 7H9 broth (BD Biosciences, San Jose, CA, USA) supplemented with an oleic acid–albumin–dextrose–catalase mixture (BD Biosciences) and mycobactin J (Allied Monitor, Fayette, MO, USA) at 37°C with constant shaking at 200 rpm for 60 d. *M. smegmatis* mc2155 cells were cultured under the same conditions but without mycobactin and for 7 d. *Escherichia coli* BL21 (DE3) was cultured in Luria–Bertani (LB) broth at 37°C for 24 h with shaking. Stock *M. smegmatis* and *E. coli* cells suspensions were adjusted to an optical density at 600 nm (OD_600_; measured using a UV-visible spectrophotometer, Cytation 5, Biotek Inc.) measurement of 1. Then four serial 10-fold dilutions of the stock suspension were made for CFU counting on LB agar plates.

### 2.2. Magnetic beads assay

Tosyl-activated Dynabeads™ M-280 (Invitrogen)(37) were used for cell capture. Briefly, 21 μl of beads was transferred to one sterile microcentrifuge tube of 2-ml capacity (microtube) and washed twice with 1 ml of PBS (pH 7.4) for 1 min each time. The microtube was placed on a magnet (Magna grip rack, Millipore) for removing supernatant. Washed beads were then added to microtubes containing 1 mL of PBS, milk, urine, and feces suspensions spiked with various levels of *M. smegmatis* or MAP cells. These tubes were placed in a microtube rotator (Thermo Scientific™ Labquake™, Edmonton, AB, Canada) at room temperature for 1 h. The microtubes were then placed on a magnetic rack for 1 min to carefully remove the supernatant to another microtube. Magnetic beads coupled to *M. smegmatis* or MAP were washed twice with 1 mL PBS for 1 min before moving on to the detection assays using qPCR or phage D29.

### 2.3. Recovery of viable *M. smegmatis* and *E. coli* cells by magnetic beads

#### M. smegmatis

Suspensions of *M. smegmatis* cells were prepared from a stock. It was adjusted to OD_600_ ~1 (microtube#1), and then serial 10-fold dilutions in PBS were performed (microtube#2, microtube#3, and microtube#4) to obtain different cell concentrations for assays. Determination of viable *M. smegmatis* recovered by beads and centrifugation was determined based on the number of cells isolated from the supernatant, as sedimenting cells may form clumps which can lead to an underestimation of *M. smegmatis* cells isolated on LB plates. To ensure all cells remained in the entire volume of supernatant, the experiments were optimized for 2-mL microtubes. Two aliquots (1 mL) of each microtube were transferred to two microtubes for centrifugation (control) and beads methods. Microtube#1 (control) was centrifuged at 3,000 *x g* at 4°C for 16 min, and the total volume of supernatant was cultured on LB agar plates for counting cells that were not sedimented. Another microtube#1 was assayed for magnetic capture, and the total volume of supernatant was carefully removed and transferred to one plastic cuvette for measuring turbidity via OD_600_ and for colony counts on LB-agar plates. The same procedure, except measuring turbidity, was applied for microtube#2 and microtube#3. From microtube#4, *M. smegmatis* cells recovered by beads and centrifugation were resuspended in 1 mL PBS, vortexed vigorously and plated on LB agar plates. Likewise, the total supernatant volume was plated on LB agar for CFU count. The experiment was repeated 3 times.

#### E. coli

From a stock of *E. coli* suspension (adjusted to OD_600_ ~1), 2 aliquots (1 mL each) were transferred to 2 microtubes to evaluate the concentration of cells by centrifugation (2,000 *x g*, 4°C, 16 min) and capture by magnetic beads. Recovery of cells by centrifugation (control) was done following the procedure described for the microtube#1 with *M. smegmatis*. The experiment was repeated 3 times.

### 2.4. MAP cells DNA extraction

DNA extraction was performed using a GenoLyse^®^ kit (Hain Lifescience, Nehren, BW, Germany) as per manufacturer’s instructions with some modifications. Briefly, MAP cells concentrated by centrifugation or magnetic beads were resuspended in 100 μl of A-Lys, vortexed, centrifuged at 10,000 *x g* for 1 min, and incubated for 5 min at 95°C in a water bath. Then, 100 μl of A-NB buffer was added to the lysate, vortexed, and centrifuged at 10,000 *x g* for 5 min. Supernatant was transferred to sterile 1.5-ml microtubes. Concentration of isolated DNA was determined by measuring sample absorbance at 260 nm using a NanoDrop ND-1000 spectrophotometer (NanoDrop Technologies Inc., Wilmington, DE, USA).

### 2.5. Quantitative qPCR assay

This procedure was done as per Singh et al. (36). Briefly, qPCR amplification targeting *IS900* and *F57* genes was done using Taqman specific probes and StepOnePlus Real Time PCR System (Applied Biosystems). The 10-μL PCR mixture comprised 5 μL of TaqMan^®^ Universal Master Mix II (Applied Biosystems) with uracil-N-glycosylase (UNG), 200 nM of each primer and 250 nM of fluorogenic probe specific to *IS900* and *F57*; the sequence of the primers and probes used were described elsewhere (36). For each template DNA sample, 4 μL was added in the reaction mixture. qPCR conditions were as follows: 50°C for 2 min for UNG treatment, initial denaturation at 95°C for 10 min, 40 cycles at 95°C for 15 sec, and at 60°C for 1 min. Each qPCR amplification targeting *IS900* and *F57* genes was run in triplicate with negative and positive controls. Obtained data were analyzed in the form of threshold cycle (Ct) values.

### 2.6. Detection of MAP cells by qPCR

This experiment was done to evaluate the recovery of MAP cells by beads and centrifugation through the detection of DNA of the total, viable and non-viable, cells sedimented. Efficiency of beads and centrifugation to capture and sediment cells, respectively, was determined by using two referential MAP concentrations (5 × 10^7^ CFU/ml and 5 × 10^5^ CFU/mL). Briefly, two serial 10-fold dilutions were prepared from the stock (5 × 10^7^ CFU/mL) of MAP suspension in PBS. Two aliquots (1 mL) of the stock were transferred to two microtubes to evaluate the recovery of MAP by centrifugation (control) and magnetic beads. MAP cells recovered by beads and centrifugation (3,000 *x g*, 4°C, 16 min) were analyzed by *F57* qPCR. The same procedure was performed for the second serial dilution containing 5 × 10^5^ CFU/mL. Each condition was repeated three times.

### 2.7. Capture of MAP in milk, urine and feces by beads

#### From milk and urine suspension

Three sterile microtubes containing 0.9 mL of PBS (pH 7.4), 0.9 mL of urine from an apparently healthy human or 0.9 ml of skim milk (Dairyland^®^) were spiked with 0.1 mL of 10^8^ CFU/mL MAP cells. Proteinase K (Pk) was added to the microtube containing milk and incubated at 70°C for 10 min before performing the magnetic beads assay. MAP cells recovered using beads were analyzed by qPCR. The same procedure was applied to other microtubes containing fewer MAP cells (10^6^, 10^5^, 10^4^ CFU/mL). The experiment was run in triplicate and repeated twice.

#### From feces suspension

Cow feces of a known MAP-shedding calf (41) was provided by the University of Calgary. Feces suspension was prepared by mixing 3 g of apparently healthy cattle feces and 6 ml of PBS in a 15-ml centrifuge microtube. After keeping the microtube in an upright position for 20 min, 0.9 mL of supernatant was transferred to another microtube and spiked with 0.1 mL of 10^8^ CFU/mL MAP suspended in PBS. One microtube with 0.9 mL PBS was spiked with 0.1 mL of 10^8^ CFU/mL MAP. Both microtubes were used for magnetic beads assays and analyzed by *F57* qPCR. The experiment was run in triplicate and repeated twice.

### 2.8. Quantification of MAP cells recovered by beads

MAP cell counts were assessed using qPCR based upon the single copy fragment *F57* (42). Briefly, DNA calibration curves were prepared for each experiment by preparing serial 10-fold dilutions of DNA sample obtained from MAP cell (adjusted to OD_600_ ~1). The initial DNA concentration of the first serial dilution was measured through optical absorbance at 260 nm using a NanoDrop ND 1000 spectrophotometer (NanoDrop Technologies Inc.). DNA molecule copy number (reported as equivalent CFU/mL) was determined using the formula: (ng DNA × 6,023 × 10^23^)/(4,829,781 × 10^9^ × 660)(43). The number of *F57* (equivalent CFU) detected in each MAP sediment recovered by beads and control was determined by using the respective threshold cycle (Ct) values. (36)

### 2.9. Magnetic separation of M. smegmatis from urine for mycobacteriophage D29 infection

The experimental procedure was adjusted as per infection of MTB by phage D29 (44) and doubling time (3 to 4 hours) of *M. smegmatis* (45). Two urine aliquots of 1 ml containing 2.0 × 10^5^ and 2.0 × 10^3^ CFU/mL *M. smegmatis* were transferred to two sterile microtubes. *M. smegmatis* captured by beads from urine were detected by phage D29 (44,46) with some modifications. Briefly, *M. smegmatis* cells captured by beads were resuspended in 150 μL of LB with 1 mM CaCl2 (LB-S) and infected with 50 μL of phage D29 stock at 10^8^ PFU/mL and 37°C. Extracellular phage D29 was eliminated with 0.03 M ferrous ammonium sulfate (FAS) for 10 min; then 0.5 mL of LB-CaCl2 was added to the microtube to stop FAS virucide activity. The total volume suspension was mixed with 4 mL of sterile molten agar (0.8%) and 1 mL of 10^8^ CFU/mL *M. smegmatis* suspension before pouring onto Petri dishes with LB agar. Petri dishes were incubated for 24 h at 37°C and the number of infection plaques was counted. The experiment was repeated 3 times.

### 2.10. Statistical analyses

Number of *IS900* and *F57* gene copies, determined in triplicate per each sediment recovered by u-TDbeads and control, were plotted and analyzed by the Kruskal-Wallis test (GraphPad Instat, version 8; GraphPad Prism, La Jolla, CA). *P*-values < 0.05 were considered statistically significant.

## 3. Results

### 3.1. Capture of *M. smegmatis* and *E. coli* by magnetic beads

Magnetic beads captured *M. smegmatis* cells in PBS with greater capture efficiency compared to *E. coli* cells (*P* <0.05). This is evident by visual inspection of supernatant remaining in cuvettes after capture by beads (Fig. 1). Cuvette B2 shows low remaining *M. smegmatis* cell contents, unlike cuvette A2 which contained *E. coli*, indicating that beads efficiently removed cells from suspension (Fig. 1). The number of viable *M. smegmatis* cells isolated from the supernatant of each suspension showed higher recovery of viable cells by beads (> 90%) than centrifugation. *E. coli* cells were not recovered by beads; approximately 94% of cells remained in the supernatant (Table 1).

**Fig. 1.**
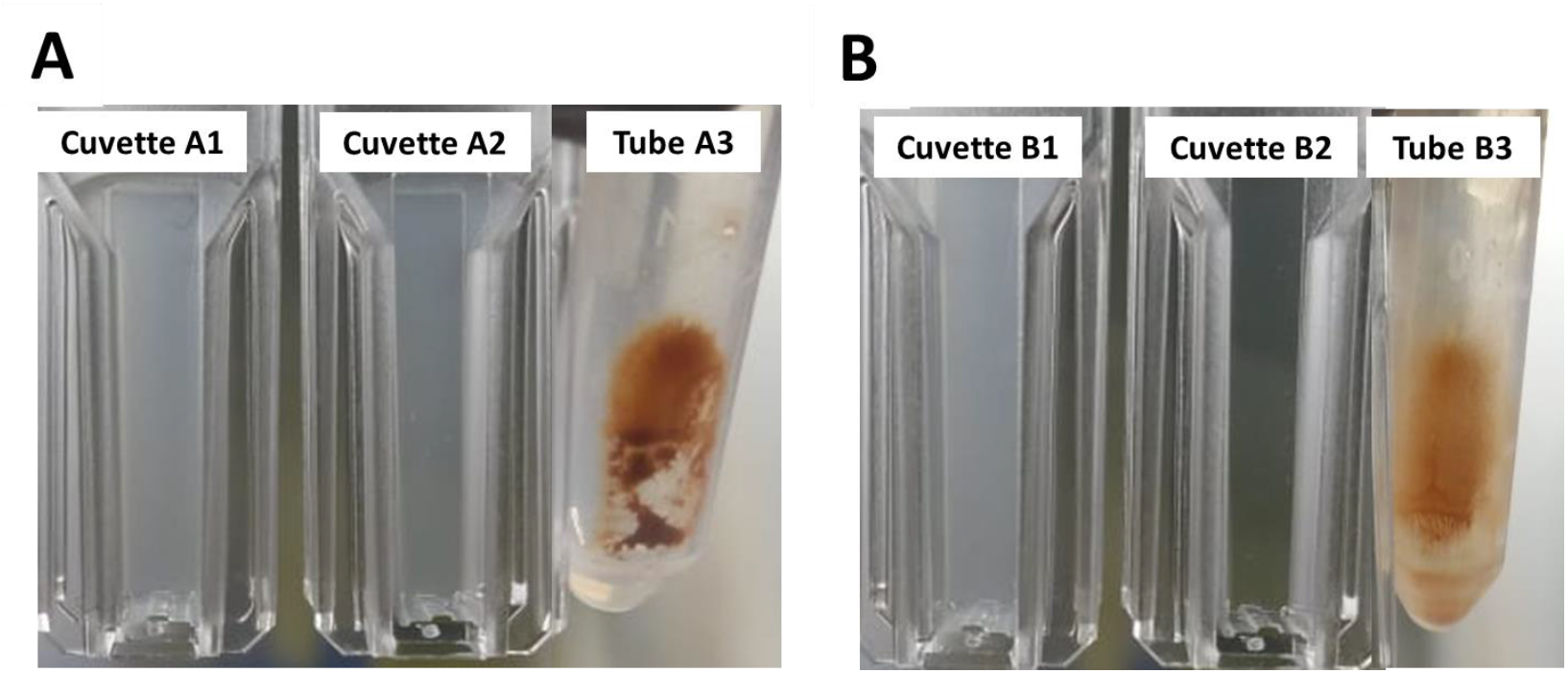
Capture of *E. coli* and *M. smegmatis* cells in PBS suspension by magnetic beads. A. Capturing *E. coli* in PBS by beads. Cuvette A1: Concentration (1,8 × 10^7^ CFU/ml) of *E. coli* before capture; cuvette A2: Concentration (1,7 × 10^7^ CFU/ml) of *E. coli* suspension after beads capture. Microtube A3: Magnetic beads bound to *E. coli* cells and separated in a magnetic rack; B. Capturing *M. smegmatis* in PBS by beads. Cuvette B1: Concentration (1,68 × 10^7^ CFU/ml) of *M. smegmatis* suspension before beads capture; cuvette B2: Concentration (6.56 × 10^5^ CFU/ml) of *M. smegmatis* suspension after beads capture; microtube B3: Magnetic beads bound to *M. smegmatis* cells and separated in a magnetic rack.

**Table 1.** *Escherichia coli* and *Mycobacterium smegmatis* remaining in supernatant after centrifugation and magnetic beads methods.

| Method | % *E. coli*<br>(mean, ±SD)<br>in supernatant<br>from (CFU/mL) | % *M. smegmatis* (mean, ±SD) in supernatant from (CFU/mL) | | | |
| --- | --- | --- | --- | --- | --- |
|  | 1,8 x 10<sup>7</sup> | 1.68 x 10<sup>7</sup> | 1.68 x 10<sup>6</sup> | 1.68 x 10<sup>5</sup> | 1.68 x 10<sup>4</sup> |
| Centrifugation | 0 | 0.66<br>(1.12x10<sup>5</sup>, ±7.64x10<sup>3</sup>) | 0 | 0 | 0 |
| Magnetic beads | 94.63<br>(1.7x10<sup>7</sup>, ±5.75x10<sup>4</sup>) | 4.04<br>(6.79x10<sup>5</sup>, ±2.16x10<sup>4</sup>) | 3.07<br>(5.17x10<sup>4</sup>, ±2.96x10<sup>3</sup>) | 6.71<br>(1.13x10<sup>4</sup>, ±9.57x10<sup>2</sup>) | 3.69<br>(6.21x10<sup>2</sup>, ±1.22x10<sup>2</sup>) |

\*Centrifugation at 3,000 x g at 4°C for 16 min

The effect of the centrifugation and magnetic beads methods on the viability of *M. smegmatis* was confirmed by assessing suspensions at 1.68 × 10^4^ CFU/mL (Fig. 2). As per one experiment results, 17 CFU of *M. smegmatis* isolated from the supernatant and 337 CFU recovered from sedimented beads, 337 CFU compressed 1326 CFU of *M. smegmatis* as consequence of the formation of clumps during the magnetic separation (Fig. 2). Thus, the magnetic beads recovered more viable *M. smegmatis* from the suspension (96%) than the centrifugation (0%). An electron microscopy image of the capture of *M. smegmatis* cells by beads is presented in Fig. 3.

**Fig. 2.**
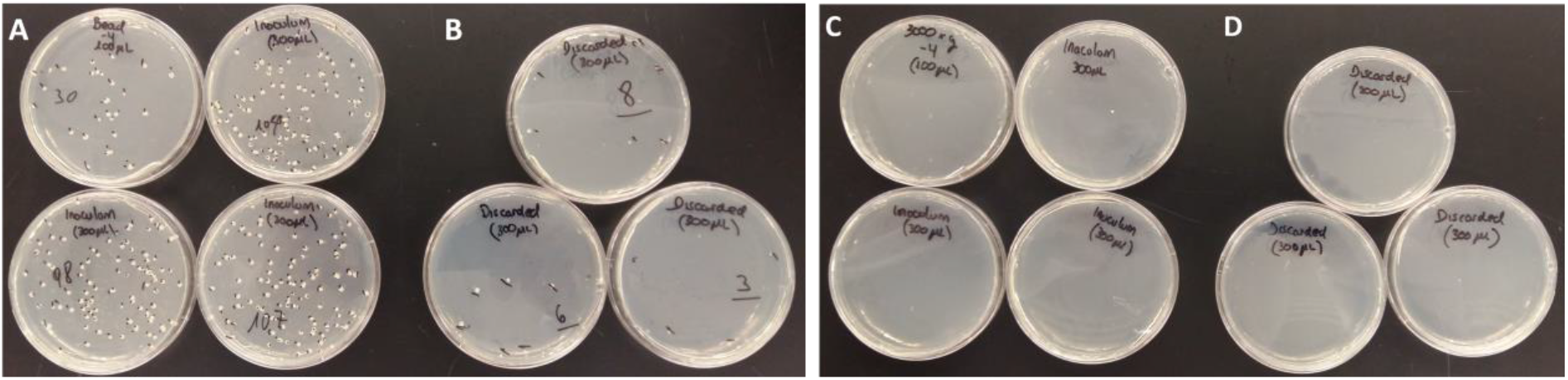
*M. smegmatis* cells recovered by magnetic beads and centrifugation. From a suspension of 1.68×10^4^ CFU/mL of *M. smegmatis*, A and B show cells captured (precipitated) and not captured (supernatant) by beads, respectively. As per the *M. smegmatis* cells remaining in the supernatant (3.7%), the estimated number of cells recovered from the beads was 96.3%. C and D show that no viable cells were recovered from the supernatant or pellet after centrifugation.

**Fig. 3.**
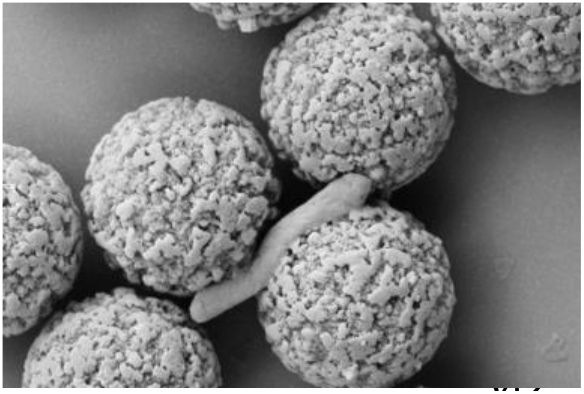
A scanning electron microscope image of a M. smegmatis cell captured by magnetic beads. Capturing *M. smegmatis* cell in PBS by magnetic beads.

### 3.2. Detection of MAP cells by qPCR

The initial concentration of MAP cells in the first and second PBS suspensions (2.93 × 10^9^ and 1.43 × 10^7^ equivalent CFU/mL, respectively) was estimated by detecting *F57* sequence from all MAP cells collected by centrifugation, which is used to concentrate 95% mycobacteria (18). Using *F57*-qPCR, there was a difference (p=0.04) in recovery of MAP cells between the magnetic beads and centrifugation methods: from the first suspension 2.8 × 10^9^ CFU/mL (Ct = 17.89) were recovered by magnetic beads and 3.2 × 10^9^ CFU/mL (Ct 17.68) by centrifugation, whereas 3.9 × 10^7^ CFU/mL (Ct = 23.94) were recovered by magnetic beads and 1.4 × 10^7^ CFU/mL (Ct = 25.43) by centrifugation ((p<0.0001); Table 2).

**Table 2.** Recovery and detection of MAP cells by magnetic beads and *F57*-qPCR.

| PBS<br>suspension | OD<sub>600</sub>, 1.0<br>(theoretical)<sup>a</sup><br>CFU/mL (±SD) | Initial MAP<br>cells<br>concentratio<br>n CFU/mL<sup>b</sup> | MAP cells recovered by<br>centrifugation* and<br>quantified by *F57*-qPCR |  | MAP cells recovered by<br>beads and quantified<br>by *F57*-qPCR |  |
| --- | --- | --- | --- | --- | --- | --- |
|  |  |  | CFU/mL (±SD) | Ct | CFU/mL (±SD) | Ct |
| **First** | 3.17x10<sup>7</sup> (±2.02x10<sup>7</sup>) | 2.93x10<sup>9</sup> | 2.8x10<sup>9</sup> (±4.1x10<sup>8</sup>) | 17.89<sup>c</sup> | 3.2x10<sup>9</sup> (±4.5x10<sup>8</sup>) | 17.68<sup>c</sup> |
| **Second** | 3.17x10<sup>5</sup> (±2.02x10<sup>5</sup>) | 1.43x10<sup>7</sup> | 1.4x10<sup>7</sup> (±1.8x10<sup>6</sup>) | 25.43<sup>d</sup> | 3.9x10<sup>7</sup> (±5.5x10<sup>6</sup>) | 23.94<sup>d</sup> |

<sup>a</sup> Approximation of CFU in each suspension was assessed by OD<sub>600</sub> measurements.
<sup>b</sup> Initial concentration (100%) of MAP cells in the first and second PBS suspensions was estimated by detecting *F57* sequence from MAP cells recovered by centrifugation (in theory 95%)\*.
<sup>c</sup> *F57*-qPCR showed that there was a statistically significant difference (p=0.04) in the recovery of MAP cells by magnetic beads and centrifugation.
<sup>d</sup> *F57*-qPCR showed that there was a statistically significant difference (p<0.0001) in the recovery of MAP cells by magnetic beads and centrifugation.

### 3.3. Milk, urine, and feces suspension

Ct values demonstrated that magnetic beads captured no less than 10^9^ CFU/mL from feces and no less than 10^4^ CFU/mL of MAP cells from milk and urine suspensions (Fig. 4). From milk suspensions, the recovery rates, confirmed by *F57*-qPCR, were 6.11% of 8.92 × 10^9^ CFUL/mL, 5.96% of 6.25 × 10^7^ CFU/mL, and 0.62% of 3.80 × 10^4^ CFU/mL. Pk was incorporated in milk suspensions as preliminary experiments showed that its activity increased the capture of MAP cells by beads (data not shown). Recovery rates of MAP cells from urine suspensions using *F57*-qPCR was 30.93% of 8.92 × 10^9^, 43.25% of 6.25 × 10^7^, 44.56% of 6.11 × 10^6^ CFU/mL, and 8.60% of 4.92 × 10^5^ CFU/mL. *IS900*-qPCR, unlike the *F57* method, was more sensitive to detect the lowest (10^4^ CFU/mL) MAP cells concentration from milk and urine (Fig. 4 and Table 3). From feces, 0.66% of 8.92 × 10^9^ CFU/mL MAP cells were captured and detected by beads and *F57*-qPCR (*IS900* was not done). MAP cells in feces suspensions at lower concentrations were not detected.

**Fig. 4.**
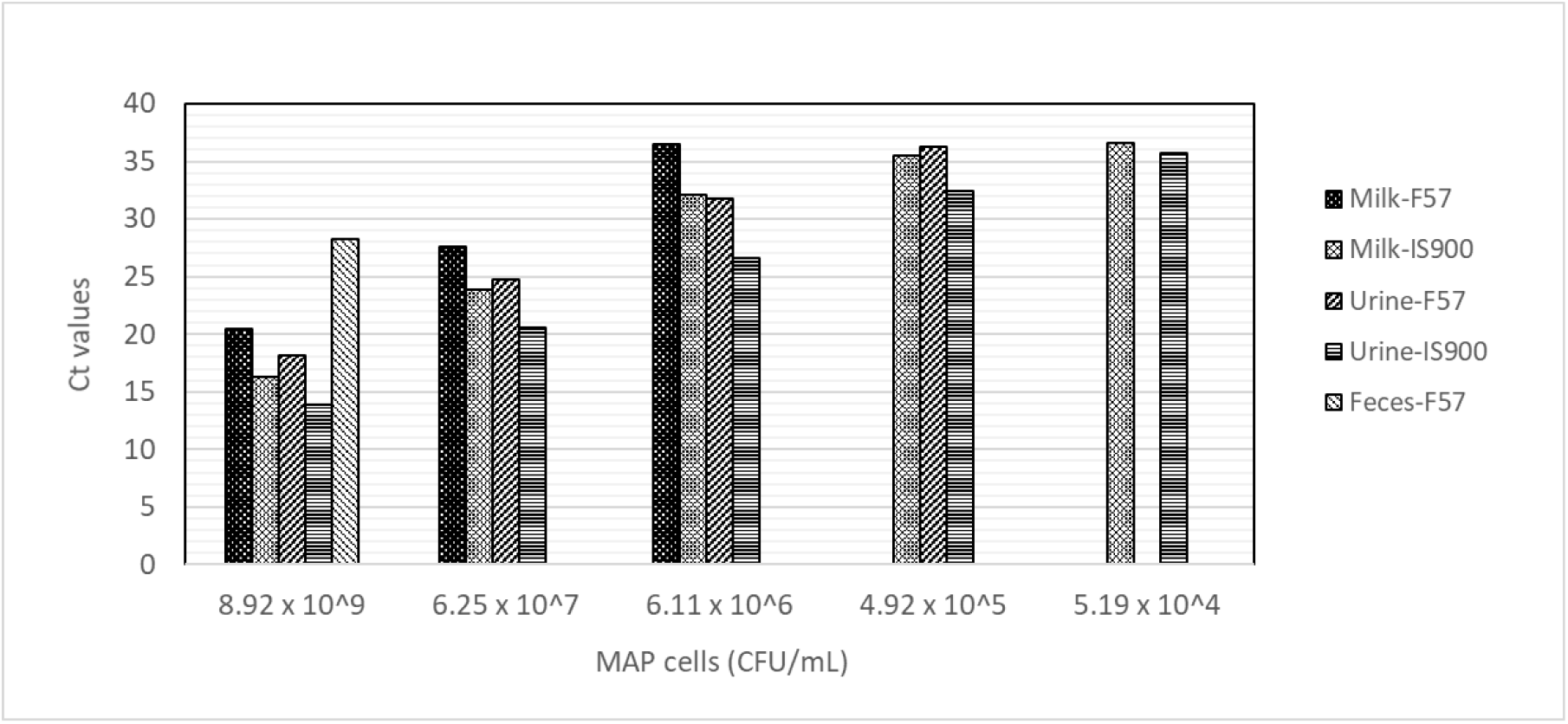
Detection of the F57 sequence by qPCR from MAP cells recovered by magnetic beads. Ct values showed that magnetic beads captured MAP cells from milk, urine, and feces. From feces, milk, and urine, *F57* sequences were detected up to 8.92 × 10^9^, 6.11 × 10^6^, and 4.92 × 10^5^ CFU/mL, respectively. *IS900*, unlike *F57*, showed better sensitivity to detect lowest (10^4^ CFU/mL) MAP cells concentration from milk and urine. Other feces suspensions containing lower concentrations were not detected by qPCR. Detection of IS sequence from feces sample was not done.

**Table 3.**
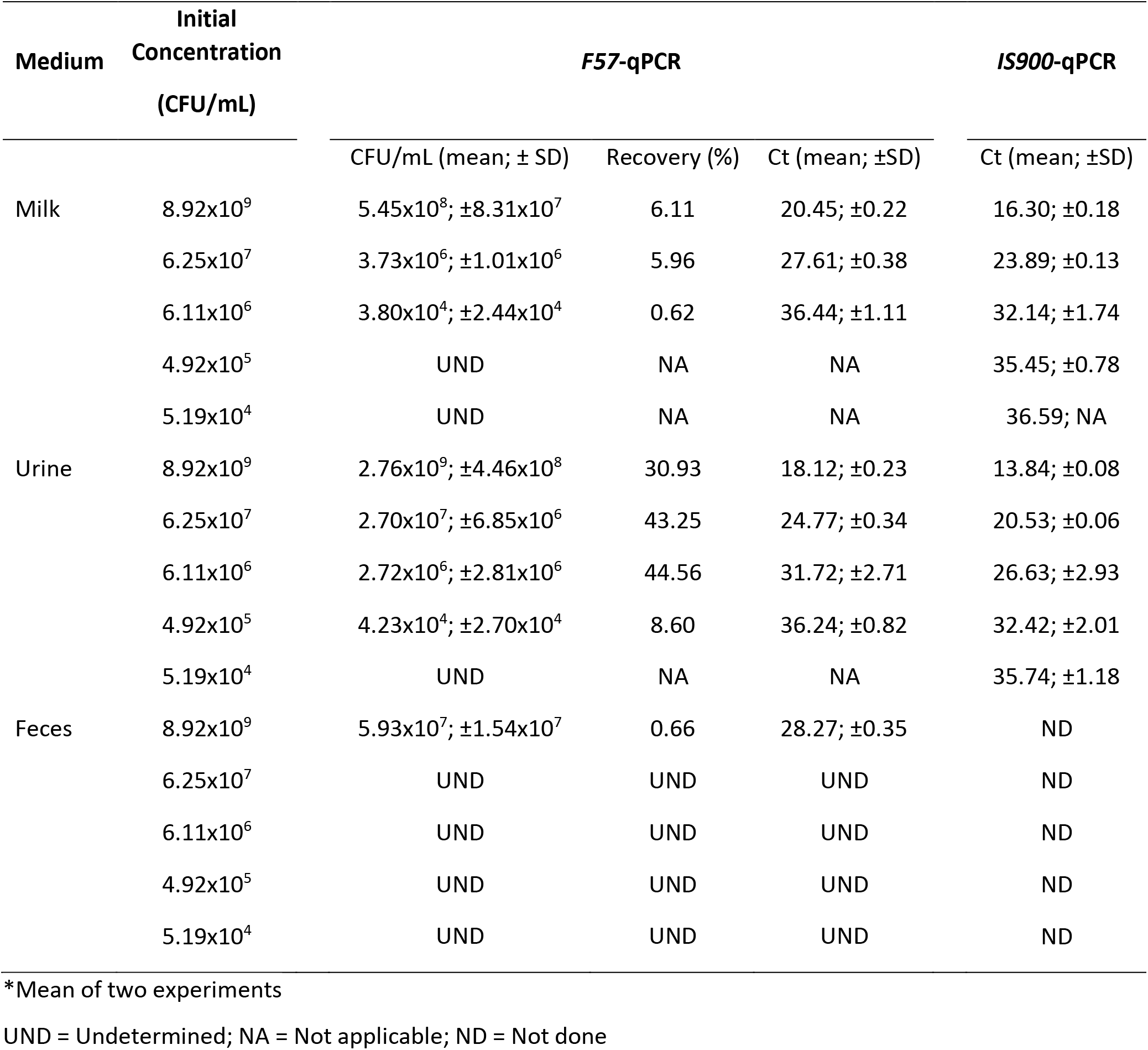
Recovery rate of *M. avium paratuberculosis* from milk, urine, and feces sample.

### 3.4. Magnetic separation of M. smegmatis from urine for mycobacteriophage D29 infection

The combination of magnetic beads and phage D29 enabled the detection of *M. smegmatis* spiked in urine within 24 h (Fig. 5). Although the number of plaques observed for samples from urine spiked with 2 × 10^5^ CFU/mL and 2 × 10^3^ CFU/mL *M. smegmatis* was lower than the respective cell counts (Table 4), the PFU detected from the samples at higher cell concentration clearly prove the detection of *M. smegmatis* within 24 h.

**Fig. 5.**
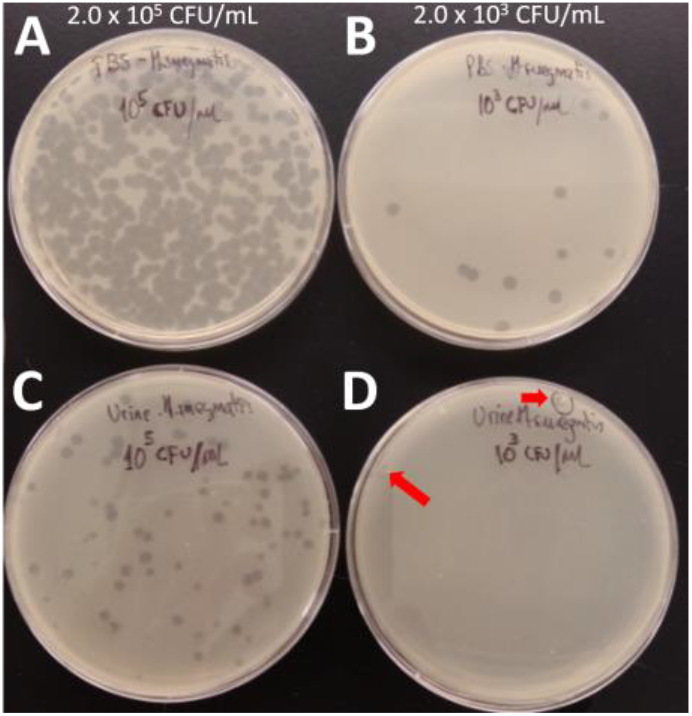
Detection of M. smegmatis from urine suspensions by mycobacteriophage D29. Detection of *M. smegmatis* captured from PBS suspensions (A and B) and urine suspensions (C and D) by mycobacteriophage D29 in 24 hours.

**Table 4.**
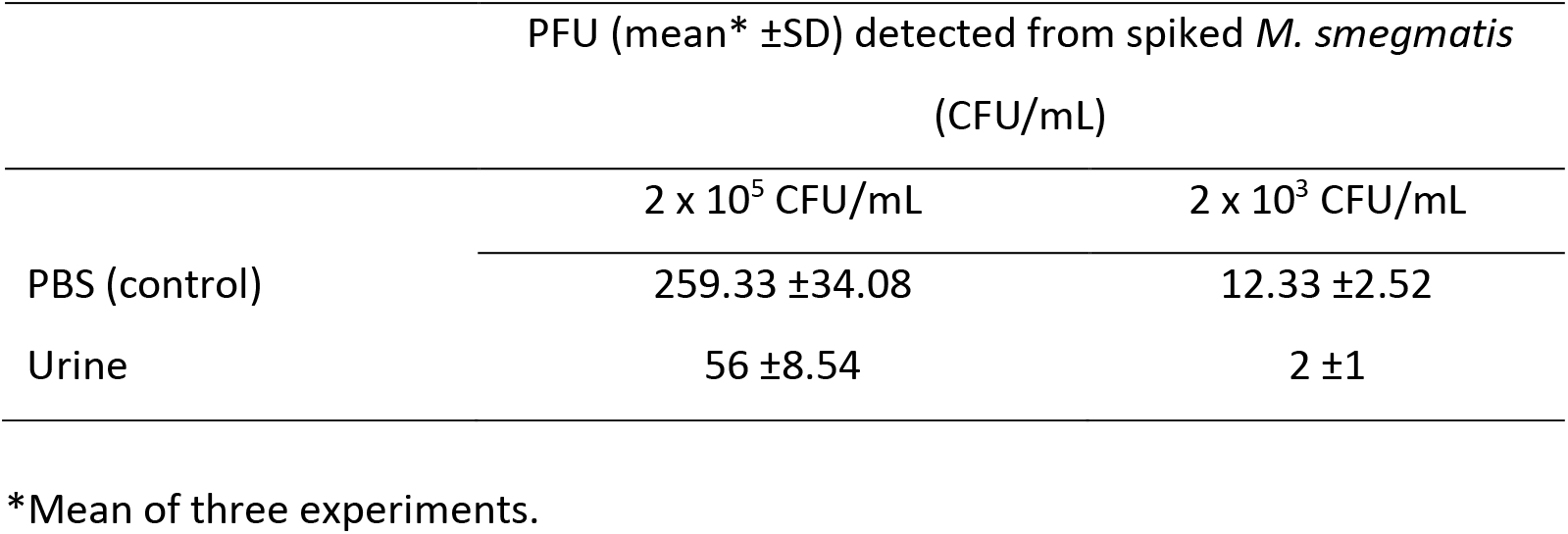
Detection of *M. smegmatis* from PBS and urine by beads and phage D29.

## 4. Discussion

Untargeted Tosyl-activated Dynabeads™ M-280 recovered more viable *M. smegmatis* cells (>90%) from spiked PBS suspension with 10^7^, 10^6^, 10^5^ and 10^4^ CFU/mL than the typical centrifuge-based method. Capture efficiency was estimated by counting unbound *M. smegmatis* cells (in supernatant) that were not captured nor sedimented by beads. Counting cells sedimented by beads, even after resuspending them in 1 mL before plating (47), could underestimate the real number of cells recovered by beads due to formation of cell clumps during the sedimentation process. With regards to cell viability, untargeted magnetic beads did not affect the viability of *M. smegmatis,* supported by growth on LB agar plate and infection by phage D29. On the other hand, centrifugation had a negative effect on the viability of *M. smegmatis* cells, regardless of cell concentration. Moreover, qPCR detection of MAP cells, after sedimentation by centrifugation and magnetic beads, indicates that the use of the latter would be preferable to precipitate cells from PBS suspension. Although the centrifugation method (3,000 *x g* at 4°C for 16 min) has been established as a referential method for the recovery of 95% of cells from suspension (18,48), we agree with others (49–51) that centrifugation may not be an appropriate method to concentrate specimens containing small numbers of microorganisms. The inefficiency of the centrifuge method may be related to the physiology of the bacteria; low recovery of MAP cells from a stationary culture is expected as the majority of cells have a lower buoyant density due to the presence of lipid bodies (52). Likewise, as others have reported for other bacteria undergoing centrifugation (53), the low number of *M. smegmatis* colonies growing on LB agar may be explained by the damage of some bacterial cells, including DNA, during the centrifugation.

Since detection of MAP cells from milk improved after treating samples with Pk, we postulate that Tosyl groups can be blocked by milk proteins which would, therefore, affect the capture of MAP cells. The positive effect of Pk on capture of MAP from milk was not observed in urine nor feces, perhaps, due to the higher protein concentration (1.87%) of milk (54) compared to urine (55) and feces. With regards to the magnetic separation of captured MAP cells from urine and feces, we postulate that the differences observed in qPCR sensitivity when detecting MAP cells in milk, urine and feces can be explained by the number of copies of the targeted genes *F57* (1 per genome) (56,57) and *IS900* (14-20 per genome)(57,58). Urea amine group, and other inhibitors such as polysaccharides originated from vegetable material in the diet (59), gut flora, bile salts, and glucolipids (60), may react with Tosyl groups and could explain decreases in the capture of cells by beads. Likewise, these inhibitors could be transferred with the beads coupled to MAP cells and reduce the amplification efficiency of *F57* and *IS900* sequences during PCR.

Magnetic beads combined with phage D29 infection detected viable *M. smegmatis* from urine suspension containing at least 10^3^ CFU/mL. Since our experiments have shown that centrifugation can affect the viability of *M. smegmatis* cells from a suspension with 10^6^ CFU/mL to the point where no viable cells remained, and considering that the decontamination of samples may damage the mycobacteria and reduce the sensitivity of phage-based assays for detecting mycobacteria cells (61), our method based on magnetic bead-mediated capture followed by phage D29 infection shows potential to detect low amounts of MAP cells from cow’s milk, and even MTB from urine samples, within 24 h. However, since the number of plaques (02 PFU) obtained from urine suspension spiked with 2 × 10^3^ CFU/mL of *M. smegmatis* may raise questions on reliability, the addition of an incubation step, enabling growth of recovered *M. smegmatis* at 37°C for 24 h, prior to phage addition could improve sensitivity by increasing the number of cells infected by phages, and by extension the number of plaques on the indicator plate. Despite that the final concentration of phages in PBS and urine suspensions (2.5 × 10^7^ PFU/mL) should ensure an infection rate of 1% (maximum infection rate) (44), the presence of beads in PBS and of urea in urine would explain the low infection rate (<1%) of cells by phages. Beads may interfere with the interaction between cells and phages, while the amino group of urea may interact with Tosyl groups and reduce the number of cells captured by beads.

Another variable that positively contributed to the capture of mycobacteria by beads was related to the technical steps required to take aliquots of beads in stock suspension, remove the supernatant after capturing beads coupled to mycobacteria and washing beads. Before taking an aliquot of the stock beads suspension, it was homogenized by pipetting 3 – 4 times carefully. The supernatant was removed without taking out the microtube from the magnetic rack to avoid removing beads coupled to mycobacteria. PBS washing was done carefully despite the fact that strong interactions are expected between beads and mycobacteria surface proteins. PBS was added slowly and carefully along the inner wall of the microtube and not directly onto the beads. The microtube should be slightly inverted 5 times to ensure that all beads were distributed homogenously and were not making visible clumps that could carry debris present in the sample. The use of 2-mL microcentrifuge tubes with attached flat top cap ensured a) homogenous mixing of suspensions for increasing interaction between beads and cells, b) removal of supernatant while the tube was placed in the magnetic rack, and c) addition of PBS to the tube for washing cells coupled to beads.

A previous study (62) demonstrating the use of Tosyl-activated Dynabeads™ M-280 coupled to Gp10, a recombinant lysin from the mycobacteriophage L5, in combination with qPCR, did not present data about the capture efficiency of uncovered magnetic beads. Another study (47) showed that other types of magnetic beads coupled to peptides and antibodies, and complemented with phages, detected approximately 0.03% and 0.003% of 10^3^ to 10^4^ CFU/mL MAP cells from milk, respectively. However, unlike the method presented in the present study, which recovered MAP cells directly from milk, the MAP cells were sedimented from milk by centrifugation and were then transferred to one microtube containing 7H9-OADC media for magnetic separation of MAP cells. Moreover, the authors (47) showed that other types of uncoated beads recovered a moderate percentage (around 10%) of nontarget mycobacteria.

A limitation of the present study was that MAP cells recovered by beads were not cultured. Although the two log10 lower counts of MAP cells between theoretical concentration determined using OD_600_ and *F57*-qPCR can be explained by clumps present in suspension and not detected using optical density, our results were in line with another study (42) that also reported a difference of two log10 counts between colony counts on culture media and *F57*-qPCR. Counting recovered cells before qPCR could further help infer the presence of death and live cells recovered by beads and centrifugation. Another variable that could not be controlled in this study was the effect of calcium as chelating agent in milk. Removing calcium from milk could reduce the formation of MAP clumps and keep a homogenous distribution of cells in suspension to be captured by beads. This study did not evaluate the effect of less amount of feces (1 g (19,63), 2g (64,65)) on the capture of MAP cells; even though this could reduce the presence and impact of inhibitors. The evaluation of these variables, and the use of target magnetic beads, in further studies would further define the potential to use untargeted magnetic beads to capture MAP cells from milk and feces.

The results of this study encourage further effort to evaluate the utility of Dynabeads^®^ M-280 Tosylactivated, with and without specific ligand, in real animals’ samples with a presumptive diagnosis of paratuberculosis. Likewise, the use of these untargeted beads, in combination with a microbiological (e.g., DS6A phage, culture media, colorimetric method) or a molecular test (e.g., HAIN-MTBDRplus, HAIN-MTBDRsl, GeneXpert), would have high potential to be used as a simple and rapid tool for diagnosis of TB.

## Author contributions

GRP designed the study and conducted the experiments. GRP, DS and RZ led the data analysis and interpretation with input from all authors. GRP wrote the manuscript and all authors edited the manuscript.

## Acknowledgment

We thank Dr. Remko van den Hurk for the scanning electron microscope image.

